# Characterizing a century of genetic diversity and contemporary antigenic diversity of N1 neuraminidase in IAV from North American swine

**DOI:** 10.1101/2022.11.18.517097

**Authors:** David E. Hufnagel, Katharine M. Young, Zebulun Arendsee, L. Claire Gay, C. Joaquin Caceres, Daniela S. Rajão, Daniel R. Perez, Amy L. Vincent Baker, Tavis K. Anderson

## Abstract

Influenza A viruses (IAV) of the H1N1 classical swine lineage became endemic in North American swine following the 1918 pandemic. Additional human-to-swine transmission events after 1918, and a spillover of H1 viruses from wild birds in Europe, potentiated a rapid increase in genomic diversity via reassortment between introductions and the endemic classical swine lineage. To determine mechanisms affecting reassortment and evolution, we conducted a phylogenetic analysis of N1 and paired HA swine IAV genes in North America between 1930 and 2020. We described fourteen N1 clades within the N1 Eurasian avian lineage (including the N1 pandemic clade) and the N1 classical swine lineage. Seven N1 genetic clades had evidence for contemporary circulation. To assess antigenic drift associated with N1 genetic diversity, we generated a panel of representative swine N1 antisera and quantified the antigenic distance between wild-type viruses using enzyme-linked lectin assays and antigenic cartography. Within the N1 lineage, antigenic similarity was variable and reflected shared evolutionary history. Sustained circulation and evolution of N1 genes in swine had resulted in significant antigenic distance between the N1 pandemic clade and classical swine lineage. We also observed a significant increase in the rate of evolution in the N1 pandemic clade relative to the classical lineage. Between 2010 and 2020, N1 clades and N1-HA pairings fluctuated in detection frequency across North America, with hotspots of diversity generally appearing and disappearing within two years. We also identified frequent N1-HA reassortment events (n = 36), which were rarely sustained (n = 6) and sometimes also concomitant with the emergence of new N1 genetic clades (n = 3). These data form a baseline from which we can identify N1 clades that expand in range or genetic diversity that may impact viral phenotypes or vaccine immunity and subsequently the health of North American swine.

## 1. INTRODUCTION

Influenza A virus (IAV) in swine is a significant respiratory pathogen that impacts animal health and causes economic losses to swine production (Kasowski et al. 2011). IAV in swine also pose a zoonotic risk with nearly 500 swine to human transmission events detected between January 2010 and May 2021 (CDC 2018). Additionally, swine can be “mixing vessels” and become infected with human, avian, and swine IAV, providing an opportunity for reassortment resulting in novel strains that may have pandemic potential (Ma et al. 2008, Rajao et al. 2019, Vincent et al. 2020). Notably, the process of reassortment has been associated with prior IAV pandemics (Medina and García-Sastre 2011, Gaymard et al. 2016). The swine-origin 2009 H1N1 pandemic is an example of the consequences of evolution and emergence of novel IAV in swine hosts (Dawood et al. 2012). Transmission from humans to swine is also common and a significant driver of IAV genetic and antigenic diversity in swine (Nelson et al. 2015, Rajao et al. 2019). Consequently, there is a need to continually assess the diversity of swine IAV to determine when and where novel IAV are emerging and whether these swine IAV pose a pandemic risk.

The two surface glycoproteins of IAV, hemagglutinin (HA) and neuraminidase (NA), have critical function in virus replication and transmission between hosts. The HA works primarily in binding IAV to the host cell surface (Bouvier and Palese 2008), and the NA has a role in releasing IAV from the host cell surface (Bouvier and Palese 2008). Due to HA/NA functional balance, the two proteins may have an epistatic relationship where any mutation in the HA that changes its binding affinity may have compensatory mutations in the NA by changing its enzymatic activity to ensure viable replication and transmission (Neverov et al. 2015, Gaymard et al. 2016, Liu et al. 2022). A consequence of the biochemical relationship is that these proteins may evolve concordantly, yet very little is known about subtype N1 diversity in swine or how this varies with paired HA diversity.

There are two cocirculating lineages of swine N1 genes in North America: The Eurasian avian (EA) lineage and the classical swine (C) lineage. The classical swine lineage is the result of a human-to-swine transmission of an H1N1 virus associated with the 1918 “Spanish flu” pandemic (Worobey et al. 2014). This lineage has persisted for over 100 years in swine, and the N1 genes in North America are most commonly paired with H1 HA genes that also share the 1918 H1N1 virus as a common ancestor, e.g., the 1A.2 (beta) and 1A.3.3.3 (gamma) H1 clades (Walia et al. 2019). The EA lineage was first detected in European swine in the 1970s and share a common ancestor with avian H1N1 (Joseph et al. 2017). The EA N1 lineage was not detected in North America until the 2009 H1N1 pandemic (H1N1pdm09); this virus was a reassortant that included the EA N1 and M gene (Garten et al. 2009, Smith et al. 2009). Following the global dissemination of the H1N1pdm09 in humans (Vijaykrishna et al. 2011), humans repeatedly reintroduced it into swine (Nelson et al. 2015, Vincent et al. 2020) and the pdm09 N1 is now a distinct genetic clade nested within the EA lineage (Vijaykrishna et al. 2011, Vincent et al. 2020). These pdm09 lineage N1 proteins in North American swine are usually paired with the pdm09 H1 HA clade (Walia et al. 2019).

Other than broad categorization to evolutionary lineage, the genetic diversity within the swine subtype N1 gene has not been quantified. The absence of a detailed classification scheme, similar to those generated for N2 genes (Zeller et al. 2021) and HA genes in swine (Anderson et al. 2016, Anderson et al. 2021) limits the ability to communicate the evolutionary and phenotypic diversity of N1 genes. Since naturally acquired NA antibodies were shown to reduce infection and illness independent of HA antibodies (Murphy et al. 1972, Couch et al. 2013), and vaccines containing both HA and NA were more efficacious than those with single antigens (Clements et al. 1986, Monto et al. 2015, Wymore Brand et al. 2022), the characterization of N1 genes will improve vaccines in swine if used to incorporate targeted N1 strains. The objective of this study was to quantify the genetic and antigenic diversity of North American N1 swine IAV genes, assess lineage and clade evolutionary rates, evaluate how reassortment affects diversity, and identify spatial and temporal patterns in N1 and NA-HA pairings. In this study we created a detailed phylogenetic classification system for N1, demonstrated antigenic differences between N1 evolutionary lineages, identified geographic and temporal hotspots of genetic diversity, and demonstrated that N1 evolution was driven by preferential pairing with HA genes. Our data demonstrate a high degree of genetic and antigenic diversity of the N1 gene in swine IAV and indicate that including matched NA can improve vaccine control efforts.

## 2. MATERIALS AND METHODS

### 2.1 Data acquisition and curation

All available swine N1 sequences from North America from 1930 to September 2020 were downloaded from the Influenza Research Database (IRD) (Zhang et al. 2017). These data were curated, and we removed mixed infection samples and sequences with >= 5% ambiguous bases. Duplicate sequences were identified, and a single representative was retained. Sequences were trimmed to the open reading frame and aligned with MAFFT v7.453 (Katoh et al. 2002) within Geneious Prime v.2020.2.2 for visualization using a custom Python curation script. Two sequences caused a frame shift (A/Swine/South Dakota/A01267895/2012 and A/Swine/Quebec/1605785/2014) and were removed. Our final step was to remove sequences with lengths less than 70% of the complete NA using smof (Arendsee et al. 2018). The dataset and custom scripts are available at https://github.com/flu-crew/N1-diversity.

### 2.2 Phylogenetic analysis, clade classification, and quantifying genetic diversity

The N1 gene data were aligned using MAFFT v7.453 (Katoh et al. 2002) and a maximum likelihood tree was inferred using IQ-TREE v.2.0-rc1 with automatic model selection (Nguyen et al. 2015). Clades were defined by sharing of a common node and monophyly; statistical support (>=90 SH-like approximate likelihood ratio test (Guindon et al. 2010), and >=75 ultrafast bootstrap (Hoang et al. 2018)); and evidence of contemporary circulation (having at least one sequence on or after January 1 2019). Subsequently, the within- and between-clade average pairwise distances were calculated using MEGAcc v.10.1.8 (Kumar et al. 2018) with a goal of having within-clade mean distances below 5% and between-clade distances above 5% (following (Anderson et al. 2016). For visualization purposes, we downsampled the inferred maximum likelihood tree using smot (https://github.com/flu-crew/smot). This algorithm sampled the square root of the number of tips in each paraphyletic branch of the N1 tree. For monophyletic clades sqrt(n) tips were randomly sampled. For paraphyletic clades, such as N1.C.3, the tree was split into segments (two, in the case of N1.C.3) and sqrt(n) random samples were chosen, where n is the number of tips in each segment. Clades with evidence of onward transmission were annotated and all data and genetic clade annotated trees are at https://github.com/flu-crew/N1-diversity

### 2.3 Assessing antigenic diversity of the N1 of IAV in swine

To assess whether between-clade genetic diversity in the N1 clades was correlated with antigenic diversity, we identified representative N1 genes for each contemporary circulating clade. Swine N1 genes collected between January 1, 2018 and July 6, 2020 were extracted from our dataset, translated to amino acid using smof (Arendsee et al. 2018), and aligned with default settings in MAFFT v7.453 (Katoh et al. 2002). A maximum likelihood phylogenetic tree was inferred using IQ-TREE v2.0-rc1 (Nguyen et al. 2015), and circulating N1 clades were identified following criteria established in section 2.2. An amino acid alignment was generated for each clade, and sequences collected between January 2020 and July 2020 were used to generate a majority consensus sequence in smof (Arendsee et al. 2018). For clades with infrequent detections, we relaxed the temporal window to begin with January 2019 for N1.C.1.1 and July 2019 for N1.C.2. A pairwise distance matrix was generated for each clade and the best-matched field strain to the clade consensus was identified and selected. If there were multiple best matches, we selected the most recently collected N1 field strain (Table S2). Representative viruses were requested from the USDA influenza A virus in swine surveillance system virus repository at the National Veterinary Services Laboratories (Arendsee et al. 2021).

Recombinant IAV (rgH9N1) were generated by pairing the NA gene with an irrelevant H9 gene from A/guinea fowl/Hong Kong/WF10/99 (H9N2) (Obadan et al. 2019) on an attenuated internal gene cassette from A/turkey/Ohio/313053/2004 (H3N2) (Pena et al. 2011, Cáceres et al. 2021, Kaplan et al. 2021). Reverse genetics viruses were obtained in a coculture of human embryonic kidney (HEK) 293T and Madin-Darby canine kidney (MDCK) cells. TransIT-LT1 transfection reagent (Mirus Bio LLC, Madison, WI, USA; 18μl total per reaction) and 1μg of each plasmid (8μg total per reaction) were combined and incubated for 45 min, then used to overlay the 293T/MDCK cells overnight. The next day the transfection mixture was replaced with fresh Opti-MEM media containing 1% antibiotics (Life Technologies, Carlsbad, CA, USA) and 24 h post-transfection the media was supplemented with 1 μg/mL of tosylsulfonyl phenylalanyl chloromethyl ketone (TPCK)-treated trypsin (Worthington Biochemicals, Lakewood, NJ, USA). The rgH9N1 viruses were propagated on MDCK cells and harvested 72 hours post-inoculations. Viruses were centrifuged, aliquoted and stored at −80°C. Virus work was performed in a biosafety level 2 laboratory in accordance with University of Georgia institutional biosafety protocols (IBC protocol #2020-0018). Plasmids and viruses’ sequences were confirmed by Sanger sequencing (Psomagen, Rockville, MD, USA) and matched the expected consensus sequence. Swine antiserum to inactivated rgH9N1 were generated in three-week-old pigs obtained from a herd free of IAV and porcine reproductive and respiratory syndrome virus. Pigs were housed in a biosafety level 2 containment facility in compliance with the USDA NADC Institutional Animal Care and Use Committee. For each rgH9N1, two pigs received an intramuscular injection of 128-256 HAU UV-inactivated virus with a 20% Emulsigen-D adjuvant (MVP Laboratories, Omaha, NE, USA). Pigs were boosted 2-3 weeks after the initial vaccination with an identical dose of the rgH9N1. When serum inhibition (HI) titers were greater than 160 HAU, pigs were humanely euthanized for blood collection.

A neuraminidase inhibition enzyme-linked lectin assay (NI ELLA) was then performed as previously described in Kaplan et al. (2021). Three dimensional antigenic maps were generated in Racmacs (Wilks 2022) from the NI data (https://github.com/flu-crew/N1-diversity). The pairwise antigenic distance between each of the antigens was extracted from the map and graphed using the ggplot2 package in R v.4.0.2 (Wickham 2016).

### 2.4 Quantifying lineage- and clade-specific evolutionary rates

To estimate the evolutionary rates of N1 NA, we created alignments for the C lineage as well as each of the following major genetic clades: N1.C.2, N1.C.2.1, N1.C.3, N1.C.3.1, N1.C.3.2, N1.P. We included all data for clades with fewer than 750 genes and randomly subsampled larger clades down to 750 genes for three replicates. N1 genes from the N1.C.1.1 clade were removed as they had incongruent divergence and sampling dates (Rambaut et al. 2016) because these N1 genes were introduced to US swine through the use of a live attenuated influenza virus vaccine that included an N1 gene collected in 1999 (Sharma et al. 2020). For each N1 clade alignment, we inferred time-scaled phylogenetic trees in BEAST v1.10.4 (Drummond and Rambaut 2007). The analyses used the SRD06 substitution model, strict molecular clocks with the exception of the N1.C clade where we implemented an uncorrelated relaxed clock with lognormal distribution, and the GMRF Bayesian Skyride coalescent tree prior (Minin et al. 2008). The MCMC were set to 200,000,000 iterations with sampling every 20,000 iterations. Convergence was assessed in Tracer v1.7.1 (Rambaut et al. 2018) and a maximum clade credibility tree was generated and annotated using TreeAnnotator v1.10.4 (Drummond et al. 2002) with 10% burn-in. The time to most recent common ancestor was inferred from the nodes between clades, using the 95% higher posterior density (HPD) as the range of uncertainty.

The mean rate of substitution was compared between N1 clades, and between the C lineage and N1.P clade. We did not calculate evolutionary rates for the EA lineage as it is infrequently detected in North America. We extracted the posterior distributions of the mean substitution rates from BEAST using a custom Python script adapted from (Zeller et al. 2021). This approach subtracts the posterior distribution of the mean rate of substitution of one clade from another, and then calculates the 95% credible interval of the resulting distribution using the HPD function in the R package BayesTwin (Schwabe 2017). The null hypothesis of no significant difference between the two compared distributions can be rejected if zero lies outside of the calculated credible interval.

### 2.5 Describing the spatiotemporal distribution of N1 clades and N1-HA clade pairs

To determine when and where N1 genetic clades and N1-HA pairings were most frequently detected, we implemented a spatiotemporal analysis in SaTScan v9.6 (Kulldorff 1997, 2021). Using a relative risk (RR) metric, this approach can identify spatial clusters of circulating genetic clades, e.g., swine in IA are more likely to have detections of N1.C.3.1 genes than other clades; and spatio-temporal clustering of genetic clades, e.g., N1.P was more likely to be detected in MN, IA, and IL between 2014 and 2018 than any other place and time (Park et al. 2016). Each N1 gene and N1-HA gene pair were assigned the state centroid latitude and longitude for spatial location, and date of detection was used as the temporal window. NA-HA clade pairings associated with less than ten strains were omitted and clades within N1.C.1 (N1.C.1, N1.C.1.1, and N1.C.1.2) were aggregated to N1.C.1.X to account for infrequent detection. A Poisson probability model (Jung et al. 2010) was used to determine where and when clades and clade pairs were overrepresented relative to the rest of the US swine population during the testing period; and the multinomial model (Kulldorff 1997) was applied to address which clades are overrepresented relative to other clades at a certain place and time. To standardize the inferred risk score provided by the Poisson model, we used the swine population of each state for each year (December 2009 to December 2020) obtained from the annual USDA NASS agricultural reports (USDA-NASS 2020). The scan statistic in both models used a circular scan window that restricts the maximum radius of a cluster to cover at most 40% of the US swine population. The SaTScan settings using populations multiplied by a factor of 1/1000, the default Mote Carlo replications for all runs (n = 999), with a resultant smallest possible resulting p-value of 0.001. For visualization, we used R v.4.0.2 with the ggplot2 package (Wickham 2016).

### 2.6 Detecting N1 and HA gene reassortment and persistence

Reassortment in the N1 NA gene was identified through annotation of the N1 NA phylogenetic tree with paired HA gene clades. A reassortment event was identified when a unique N1-HA clade pairing was detected within a monophyletic clade of a different N1-HA clade pairing. We considered novel pairings to be significant when the donor HA clade was derived from a different HA monophyletic clade, e.g., a significant reassortment would be an N1 exchanging a paired H1-1A.1 to a H1-1A.3.3.3. Reassortment events from HA clades that shared a recent common ancestor, e.g., 1A.1 to 1A.1.1, were not used in subsequent analysis. Reassortment events were considered sustained when at least ten strains downstream of the reassortment event on the tree were associated with the new NA-HA clade pair.

## 3. RESULTS

### 3.1 Extensive within-lineage diversity in swine N1 genes

3,730 N1 gene sequences from North American swine from 1930 to September 2020 were analyzed, two major evolutionary lineages were detected, and 14 statistically supported genetic clades were identified (Figure 1, Table 1). The N1 genes were collected in 32 of the 50 US states and Canada, Mexico, Costa Rica, Cuba, and Guatemala (Figure 2, Table S1).

**Figure 1.**
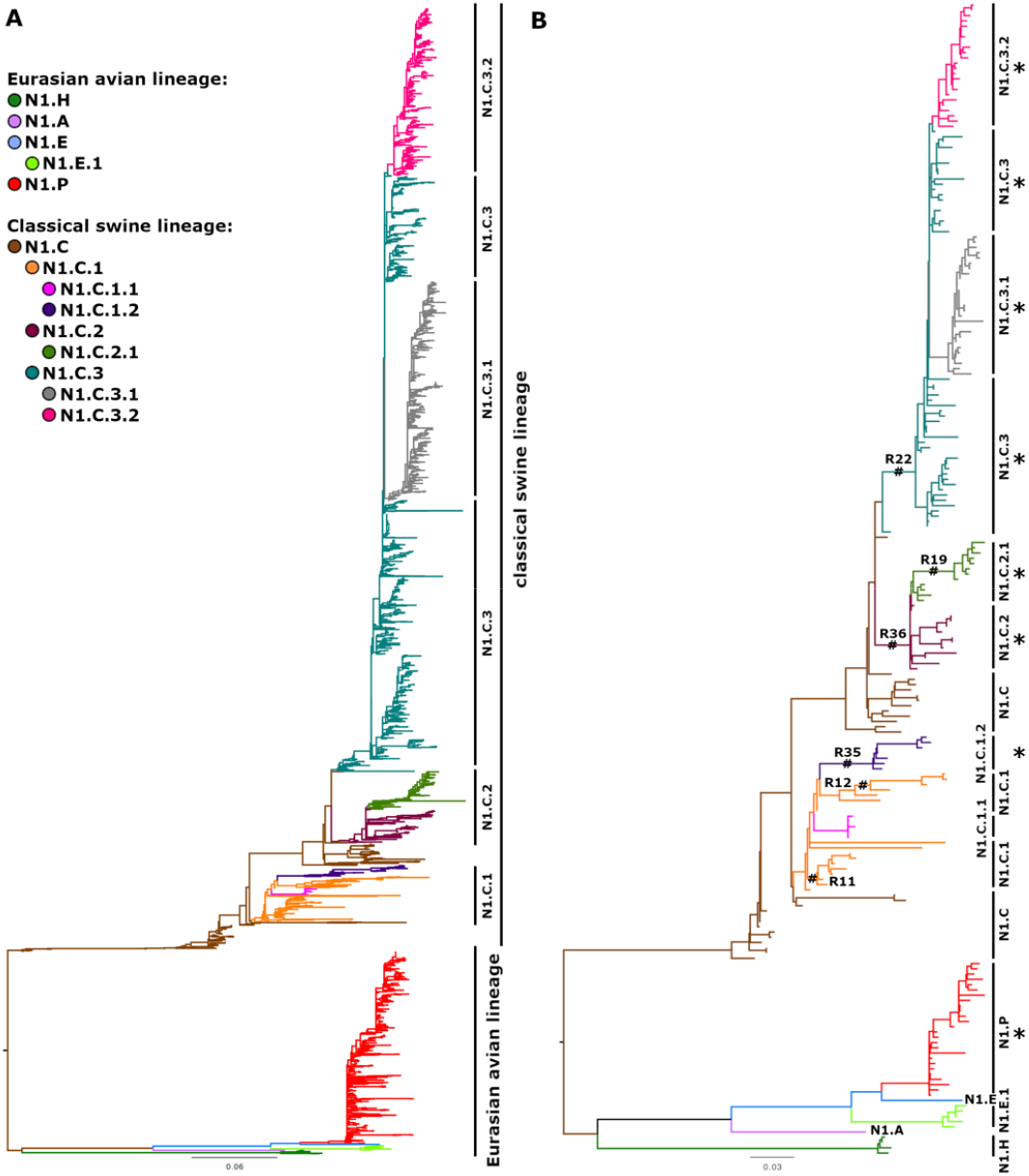
Phylogenetic relationships of N1 genes detected in North America between 1930 and 2020. (A) Phylogenetic tree with all genes following curation. (B) A sqrt(n) subsampled gene tree rooted to maintain the monophyletic lineage topology for the Eurasian avian lineage and the Classical swine lineage. Clades with evidence of contemporary circulation are indicated by asterisks. Sustained reassortment events are indicated by hash symbols and labeled by a reassortment event number.

**Table 1.**
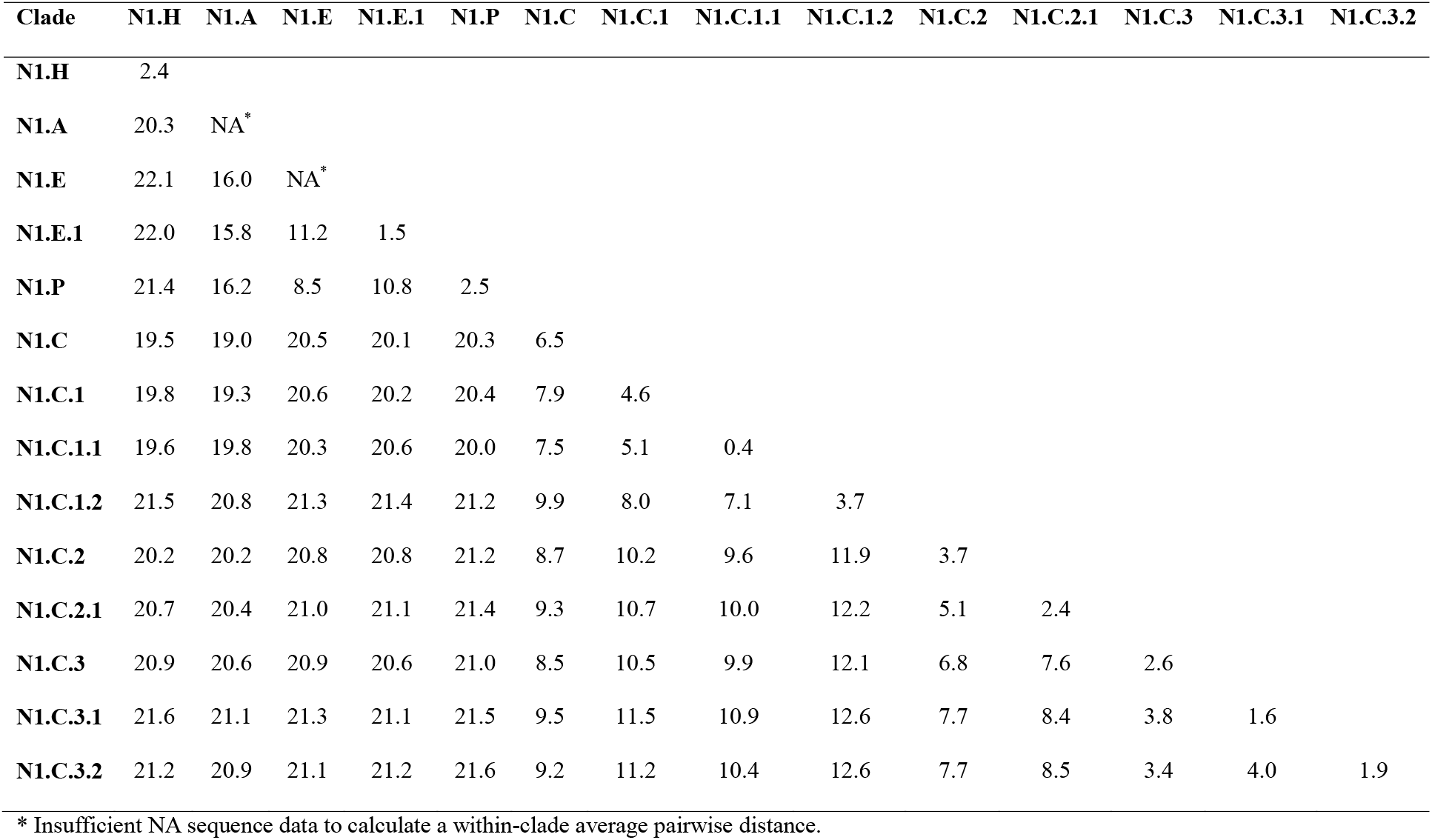
Average within- and between-clade pairwise distances for N1 neuraminidase influenza A virus in swine genetic clades. Within-clade average pairwise distance is presented on the diagonal.

| Clade | N1.H | N1.A | N1.E | N1.E.1 | N1.P | N1.C | N1.C.1 | N1.C.1.1 | N1.C.1.2 | N1.C.2 | N1.C.2.1 | N1.C.3 | N1.C.3.1 | N1.C.3.2 |
| --- | --- | --- | --- | --- | --- | --- | --- | --- | --- | --- | --- | --- | --- | --- |
| **N1.H** | 2.4 |  |  |  |  |  |  |  |  |  |  |  |  |  |
| **N1.A** | 20.3 | NA* |  |  |  |  |  |  |  |  |  |  |  |  |
| **N1.E** | 22.1 | 16.0 | NA* |  |  |  |  |  |  |  |  |  |  |  |
| **N1.E.1** | 22.0 | 15.8 | 11.2 | 1.5 |  |  |  |  |  |  |  |  |  |  |
| **N1.P** | 21.4 | 16.2 | 8.5 | 10.8 | 2.5 |  |  |  |  |  |  |  |  |  |
| **N1.C** | 19.5 | 19.0 | 20.5 | 20.1 | 20.3 | 6.5 |  |  |  |  |  |  |  |  |
| **N1.C.1** | 19.8 | 19.3 | 20.6 | 20.2 | 20.4 | 7.9 | 4.6 |  |  |  |  |  |  |  |
| **N1.C.1.1** | 19.6 | 19.8 | 20.3 | 20.6 | 20.0 | 7.5 | 5.1 | 0.4 |  |  |  |  |  |  |
| **N1.C.1.2** | 21.5 | 20.8 | 21.3 | 21.4 | 21.2 | 9.9 | 8.0 | 7.1 | 3.7 |  |  |  |  |  |
| **N1.C.2** | 20.2 | 20.2 | 20.8 | 20.8 | 21.2 | 8.7 | 10.2 | 9.6 | 11.9 | 3.7 |  |  |  |  |
| **N1.C.2.1** | 20.7 | 20.4 | 21.0 | 21.1 | 21.4 | 9.3 | 10.7 | 10.0 | 12.2 | 5.1 | 2.4 |  |  |  |
| **N1.C.3** | 20.9 | 20.6 | 20.9 | 20.6 | 21.0 | 8.5 | 10.5 | 9.9 | 12.1 | 6.8 | 7.6 | 2.6 |  |  |
| **N1.C.3.1** | 21.6 | 21.1 | 21.3 | 21.1 | 21.5 | 9.5 | 11.5 | 10.9 | 12.6 | 7.7 | 8.4 | 3.8 | 1.6 |  |
| **N1.C.3.2** | 21.2 | 20.9 | 21.1 | 21.2 | 21.6 | 9.2 | 11.2 | 10.4 | 12.6 | 7.7 | 8.5 | 3.4 | 4.0 | 1.9 |

\* Insufficient NA sequence data to calculate a within-clade average pairwise distance.

**Figure 2.**
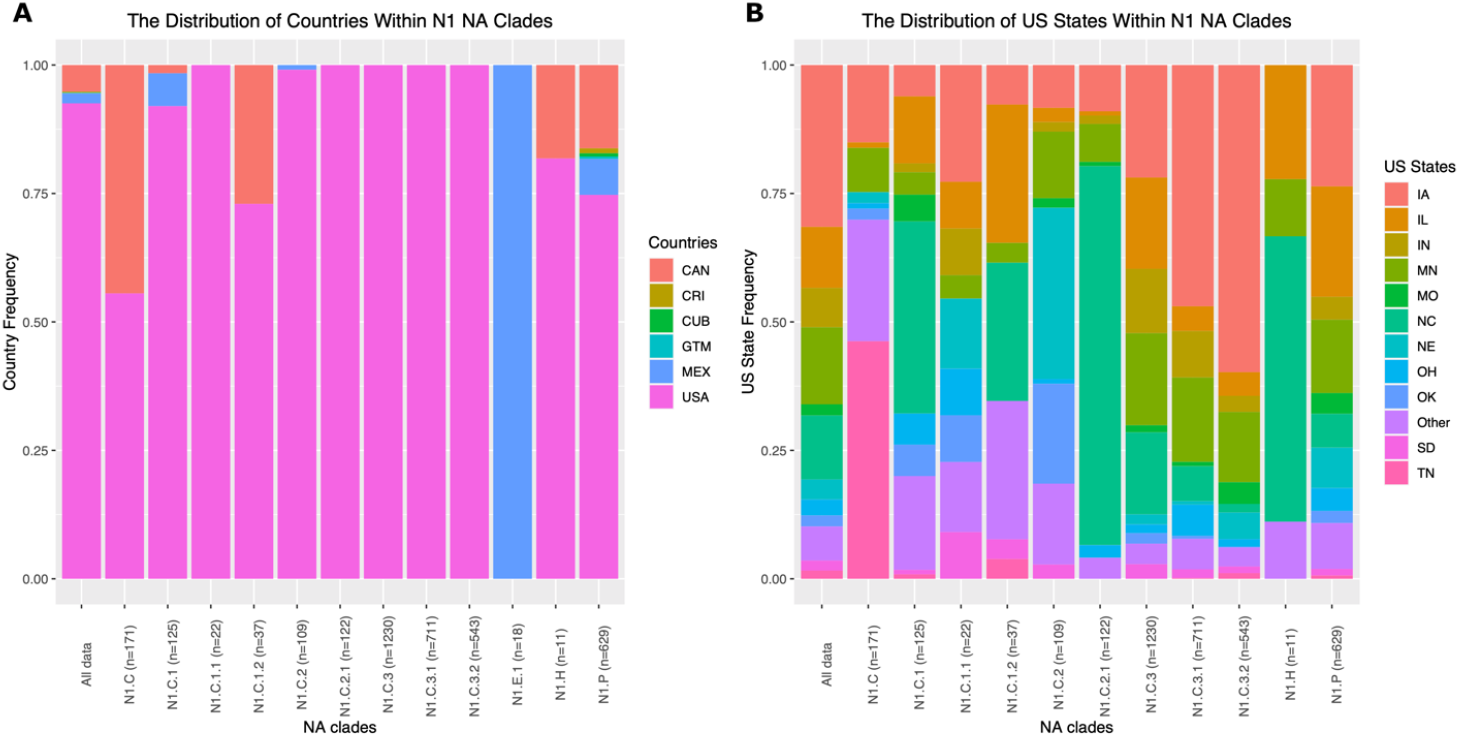
The detection proportions of N1 neuraminidase (NA) genetic clades across countries in North America (A) and within US states (B). Each vertical bar represents a statistically supported NA clade, and the color represents the fraction of samples within a given clade from each country or state. Sample sizes are provided on the *x* axis following the clade names, excluding detections where location information was not available.

There were 660 N1 genes classified within the Eurasian avian (EA) lineage across 5 statistically supported genetic clades (Figure 1, Table 1). 629 sequences from 2009 to 2020 were related to the H1N1pdm09 and classified as N1.P and this clade was detected in 22 states and 6 countries. Twelve of the remaining 31 sequences were classified based upon host origin: 11 human seasonal origin N1 genes were detected in swine from 2005 to 2009 and classified as N1.H; a single N1 gene detected in 2002 in Canada was of avian origin and classified as N1.A; The remaining EA lineage genes were detected in Mexico between 2010 and 2015 and classified as N1.E.1 or N1.E.

The classical swine (C) lineage contained 9 clades (Figure 1) consisting of 3,070 genes (82.3% of N1 detections). A first-order clade, N1.C, was introduced to classify older N1 genes with no evidence of onward circulation: this included 85 genes detected between 1930 to 2016 in 12 US states and Canada (Figure 2, Table S1). Within N1.C, three second-order divisions were identified: N1.C.1, N1.C.2, and N1.C.3. The N1.C.1 (n=184) was detected in 18 US states and 3 countries between 1987 to 2019. Third-order clades included N1.C.1.1 genes (n=22 genes, 10 US states) associated with a live attenuated influenza virus vaccine (Sharma et al. 2020), and N1.C.1.2 (n=37) detected between 2000 to 2018 across ten US states and Canada. The N1.C.2 contained 231 detections collected in 13 US states and Mexico. A monophyletic clade, N1.C.2.1, was identified with 122 sequences detected between 2007 and 2020 in 11 US states. N1.C.3 was the largest clade, encompassing 2,484 sequences collected between 2002 to 2019 across 24 US states. N1.C.3 contained N1.C.3.1 (n=711) with sequences collected between 2012 to 2020 in 22 US states and N1.C.3.2 (n= 543) with sequences collected between 2013 to 2020 in 17 US states (Figure 2, Table S1).

The within- and between-clade average pairwise distances were generally <= 5% and >= 5% respectively (Table 1). These thresholds were exceeded for the polyphyletic clade N1.C and reflected sparse sampling and independent geographic circulation of the genes across decades. We relaxed these thresholds for N1.C.3, N1.C.3.1 and N1.C.3.2 which had between-clade average pairwise distances of 3% to 4% within N1.C.3.X (Table 1), and in doing so were able to divide 2,484 genes into three groups for further analysis.

### 3.2 Significant antigenic diversity between and within N1 lineages

Antigenic cartography was used to visualize and quantify the antigenic distances between swine reference antigens (Table S2). The average antigenic distance between the representative N1.P gene (A/swine/Iowa/A02479002/2020) and C lineage genes was 6.8 AU (Figure 3). Within the C lineage there was an average of 2.0 AU between each of the N1 clades, but there was significant variation in the magnitude of the differences (range of 1.1 AU to 4.4 AU). N1.C.3.2 was over 3.1 AU from all N1 clades, except for the N1.C.3.1 clade at 0.7 AU. N1.C.1.1 and N1.C.1.2 averaged 2.2 AU and 2.4 AU away from other N1 clades respectively. The N1.C.2.X clades were generally more similar to the other representative genes, with 1.7 AU and 1.9 AU distance from N1.C.2 and N1.C.2.1 to other clades respectively. Other than the N1.C.2.X clades, each of the other representative genes had at least one pairwise antigenic distance that exceeded 3 AU. Clades that shared a more recent common ancestor (Figure 1) tended to be more antigenically similar, e.g., N1.C.3.1 and N1.C.3.2 were 1.1 AU apart.

**Figure 3.**
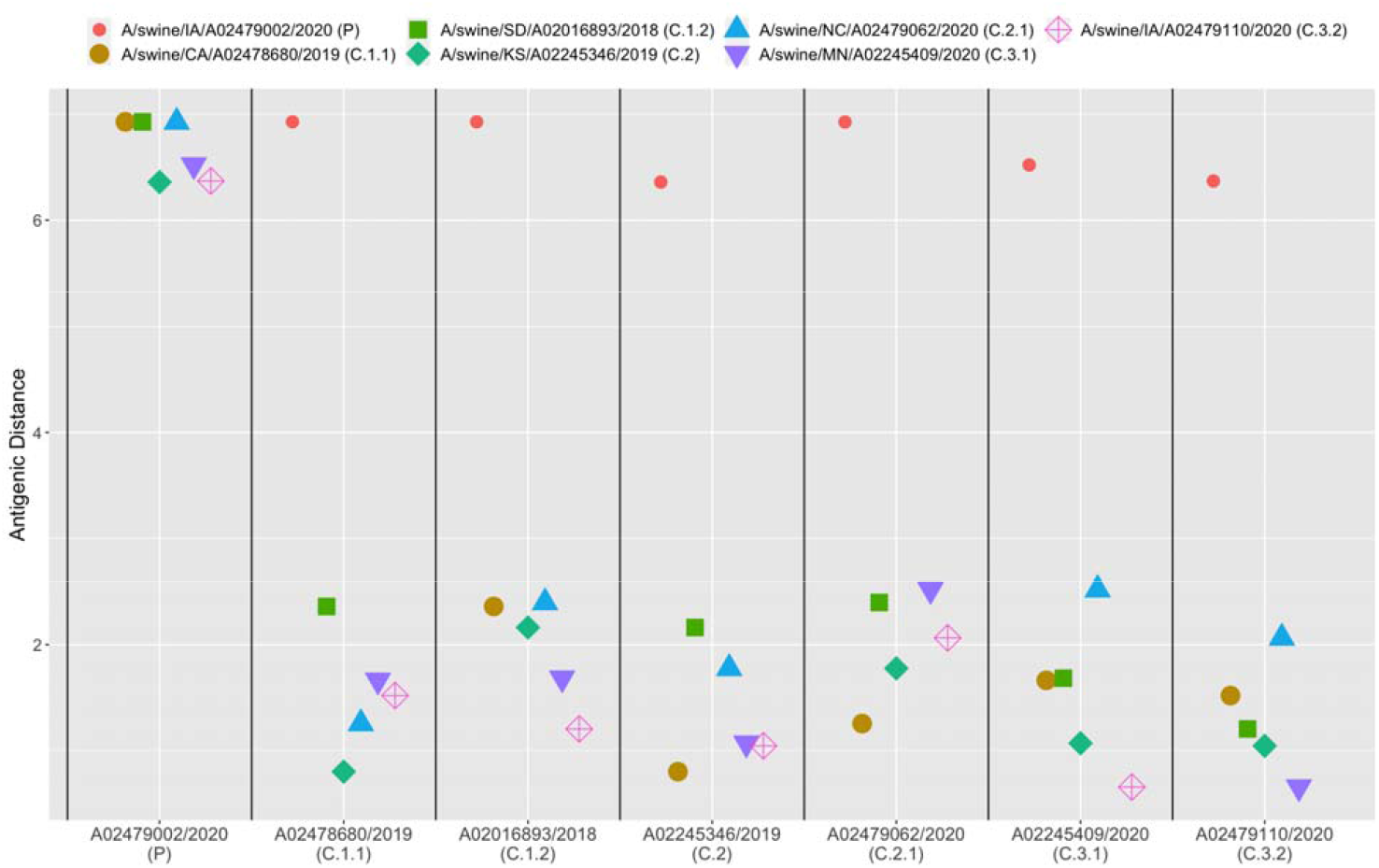
Antigenic distance between reference swine N1. The neuraminidase inhibition (NAI) of swine N1 antisera against homologous, reference antigens were determined using the neuraminidase inhibition enzyme-linked lectin assay. Antigenic cartography was used to chart the NAI data in three dimensions and calculate antigenic distances between antigens. Each antigen is listed on the *x* axis, and the pairwise distances to the other reference antigens are represented by colored shapes.

### 3.3 The Eurasian avian lineage evolved faster than the classical swine lineage

Evolutionary rates were quantified using Bayesian methods that estimated the molecular substitution rate for each N1 lineage. Given sparse sampling and detections of the EA lineage in our data, we only measured the evolutionary rate of the N1.P clade nested within the EA lineage. The N1.P clade had a substitution rate of 4.72 × 10^−3^ substitutions/site/year, and the C lineage had a substitution rate of 3.69 × 10^−3^ (Figure 4 and Table S3). The substitution rate of N1.P genes was significantly higher than that of the C lineage (Figure S1), and those of some individual C lineage clades (N1.C.2, N1.C.3.1, and N1.C.3.2). There was no statistical difference in the substitution rates between N1.C.2.1 and N1.C.3 compared to the N1.P substitution rate.

**Figure 4.**
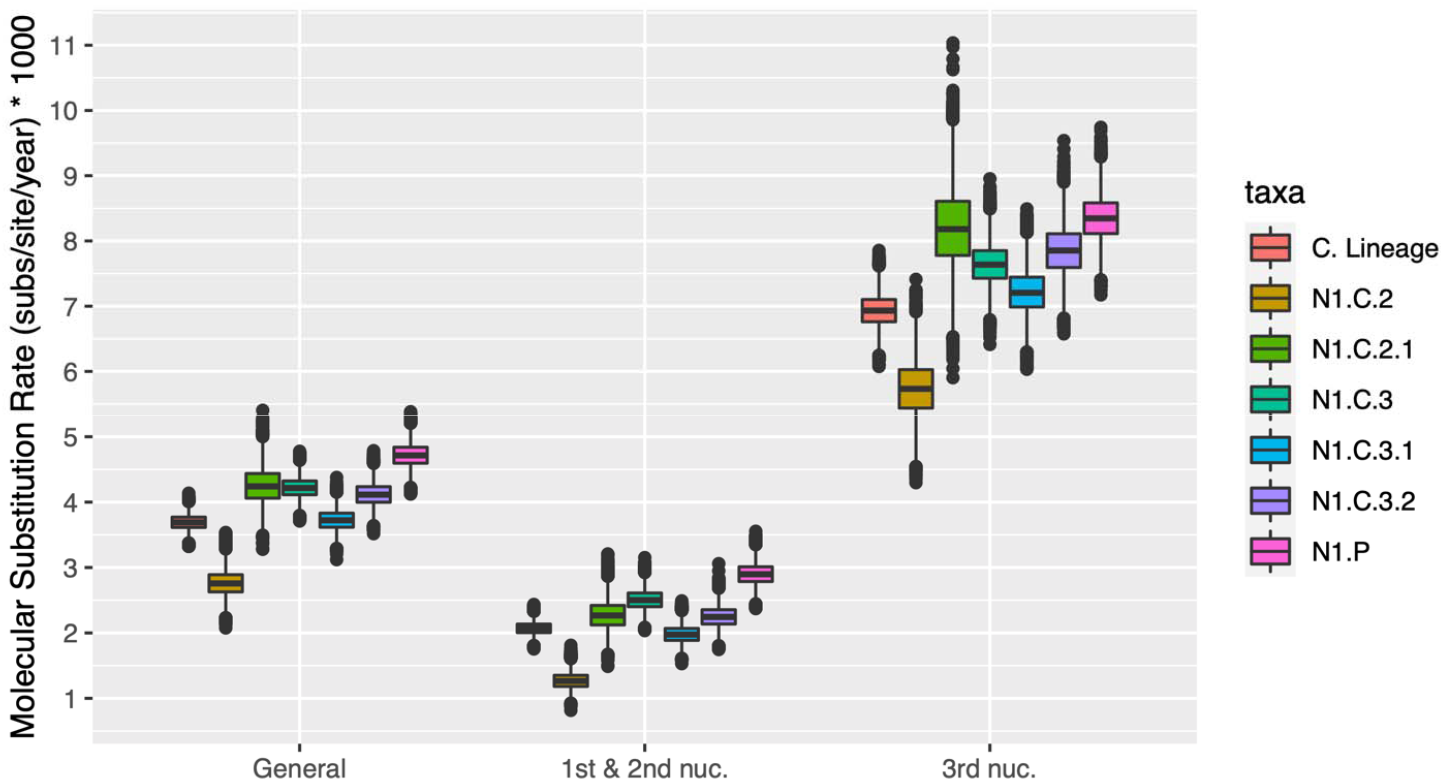
The mean nucleotide substitution rate of N1 neuraminidase clades. A mean substitution rate is presented alongside the substitution rate at the first and second codon position, and the third codon position. Rates were calculated using BEAST and implemented the SRD06 substitution model, strict molecular clocks with the exception of N1.C clade where we implemented an uncorrelated relaxed clock with lognormal distribution, and the GMRF Bayesian Skyride coalescent tree prior. The three clock rates for each codon position were averaged to determine the mean nucleotide substitution rate across the entire gene.

### 3.4 Contemporary classical swine lineage clades emerged across decades and evolved at different rates

The substitution rates and the time to the most recent common ancestor of clades with at least 50 individuals and evidence of onward circulation were quantified (Figures 3, Figure S2 and Table S4). The EA lineage did not achieve convergence, so our results were restricted to the C lineage except for the minor N1.C.1.1 clade. The N1.C.1 clade diverged from N1.C around 1984 (1980.21 – 1992.00 95% HPD), followed by N1.C.1.2 around 1991 (1989.85 – 1996.20 95% HPD). N1.C.2 and N1.C.3. and N1.C.2 and N1.C.3 diverged in approximately 2000 (1999.56 – 2003.38 95% HPD), with the second order N1.C.2.1 emerging in approximately 2006 (2006.02 – 2006.46 95% HPD). N1.C.3.1 and N1.C.3.2 originated separately in ~2012 (N1.C.3.1: 2012.00 – 2012.19 95% HPD; N1.C.3.2: 2012.29 – 2012.84 95% HPD). All N1 clades which have emerged since 2005 were detected within two years of emergence, and of the 14 named clades only N1.C.1.2 demonstrated a considerable lag from emergence in ~1991 to first detection in ~2000 (Table S4). The fastest evolving clade in our data set was N1.C.2.1 with a substitution rate of 4.26 × 10^−3^ substitutions/site/year; the slowest evolving clade was N1.C.2 with a substitution rate of 2.76 × 10^−3^ substitutions/site/year (Figure 4, and Table S3). Since diverging from N1.C.3, the evolutionary rate of N1.C.3.1 significantly decreased from 4.22 × 10^−3^ substitutions/site/year to 3.73 × 10^−3^ substitutions/site/year, but N1.C.3.2 (4.12 × 10^−3^ substitutions/site/year) did not markedly change. The substitution rate of the third nucleotide position was higher than that of the first and second nucleotides for all clades (Figure 4 and Table S3).

### 3.5 Significant variation in the spatial and temporal detection of N1 and N1-HA clade pairings

Spatial and temporal patterns associated with N1 diversity were assessed using SaTScan (Kulldorff 1997, 2021) with two probability models on N1 clade detections between January 2010 to December 2020 (Figure 5). The N1.P clade was overrepresented in Illinois between January 2010 through March 2011 (RR= 15.36, p = 0.001), in all states from November 2010 to March 2011 (RR = 4.12, p = 0.001), in Nebraska from December 2015 through June 2016 (RR = 10.85, p = 0.001), and in Iowa from April through June 2019 (RR = 9.48, p = 0.001). The N1.C.1.X clades were overrepresented in nine mostly Southern states (AR, MO, MS, LA, TN, OK, AL, and IL) from January 2011 through March 2011 (RR = 48.32, p=0.001), and in all states from March 2018 through May 2019 (RR = 5.72, p = 0.004). N1.C.2 were overrepresented in Nebraska and South Dakota from November 2011 through September 2013 (RR = 77.03, p = 0.001). N1.C.2.1 were overrepresented in N. Carolina and Virginia at all times (SG04, RR = 33.37, p = 0.001). N1.C.3 were overrepresented in Minnesota and the Dakotas from September 2012 through February 2013 (RR = 12.98, p = 0.001) and in eight states (KY, OH, IN, TN, WV, VA, NC, SC, and IL) from July 2013 through June 2015 (RR = 7.71, p = 0.001). N1.C.3.X clades tended to be overrepresented in midwestern and Great Plains states. N1.C.3.1 were overrepresented in Kentucky, Ohio, and Indiana from January 2016 through November 2017 (RR = 7.25, p = 0.001), in Minnesota from November 2017 through October 2019 (RR = 4.51, p = 0.001), and in Iowa from February 2018 through January 2020 (RR =5.23, p = 0.001). N1.C.3.2 were overrepresented in five midwestern states (NE, KS, MN, ND, and SD) from February 2015 through May 2015 (RR =12.27, p = 0.001) and in Iowa from December 2016 through November 2018 (RR =6.19, p = 0.001). The spatial and temporal clusters identified for the single N1 genes (Figure 5a) were largely mirrored for N1-HA pairs (Figure 5b).

**Figure 5.**
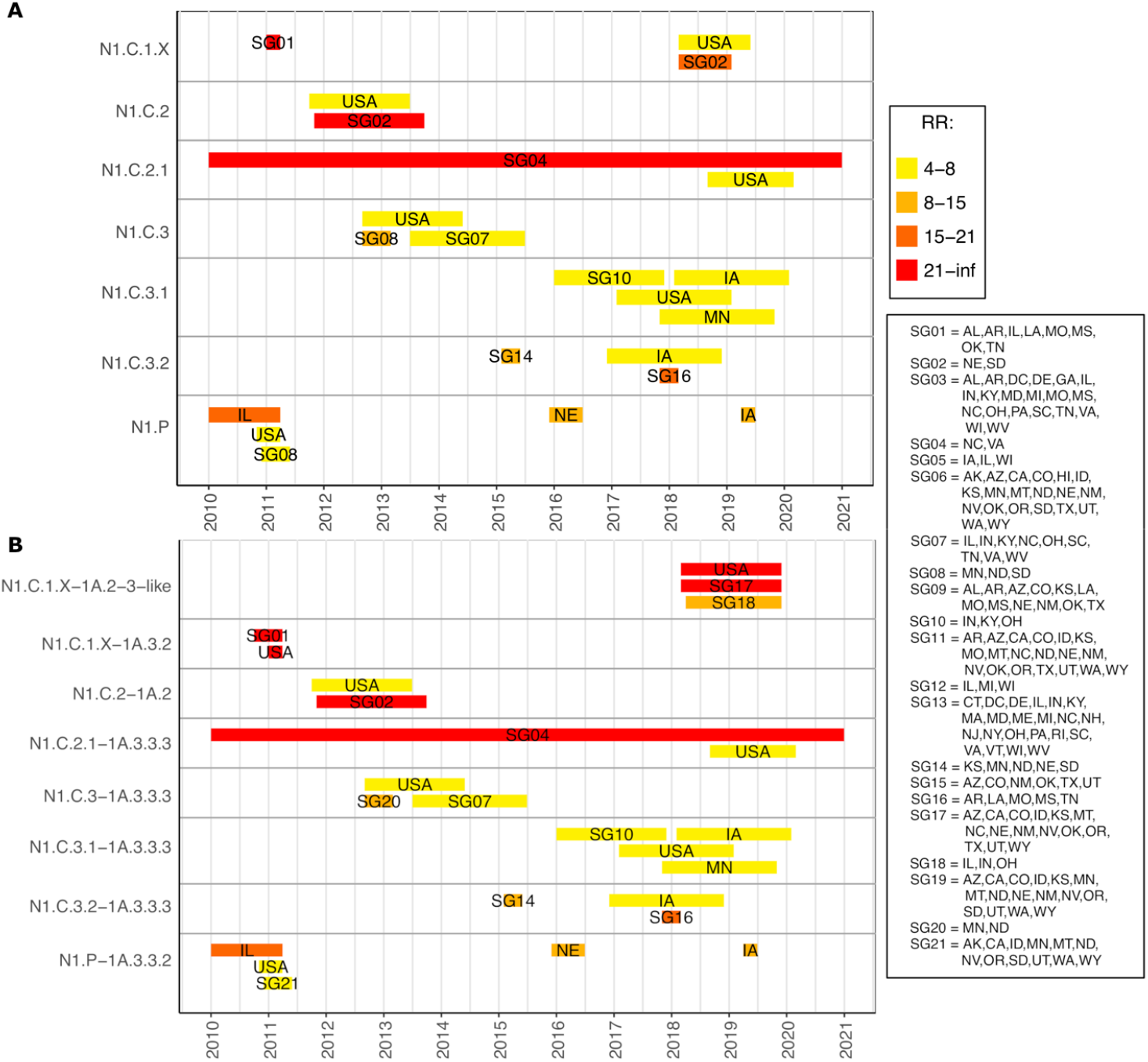
Spatiotemporal N1 clade (A) and N1-HA (B) clade pair hotspots. Spatiotemporal groups (SGs) are plotted for each clade as rectangles across a timeline from 2010 through 2020. Relative risk (RR) values from the Poisson model in SaTScan are represented by the background color of each SG rectangle. RR is the likelihood of swine in certain states at certain times being infected with a certain clade or clade pair of IAV relative to all other times and places in the USA given the swine population in each state. No RR values were exactly 8, 15, or 21. SG are groups of contiguous U.S. states that belong to a circular scan window during specific time ranges

### 3.6 Novel N1-HA gene pairings rarely sustained transmission

We identified when N1 NA genes acquired a novel HA gene pairing from 1930 to 2020. Of the 36 detected N1-HA reassortment events, only six were sustained (16.7%) (Table 2, Figure 1). The reassortment event R19 involved a N1.C.2.1 paired with a H1-1A.2 reassorting to an H1-1A.3.3.3 and 99.1% of downstream detections of N1.C.2.1 maintained the resultant H1-1A.3.3.3 pairing (106 of 107 N1-HA pairings). R22 occurred early in the history of N1.C.3 changing the pairing from N1.C.3/H1-1A.2 to N1.C.3/H1-lA.3.3.3. The new HA pairing was maintained across all subsequent N1.C.3.1 and N1.C.3.2 genes (2,429 downstream strains). Two sustained reassortment events, R35 and R36, were correlated with the formation of new N1 clades. R35 described N1.C.1 paired with H1-1A.1 HA transitioning to a N1.C.1.2 paired with an H1-1A.2. R36 described N1.C paired with a H1-1A.1.1 reassorting to a N1.C.2 paired with an H1-1A.2, which was carried through the start of N1.C.2.1 as well as the start of N1.C.3 and the one N1.C strain that sits at the base of N1.C.3 (Figure 1, & Table 2). The sustained reassortment events R11 and R12 involved N1.C.1, with the first reassorting from an H1-1A.1-like to a H1-1A.3.3.3 (71 downstream strains) and the second changing pairing from H1-1A.3.3 to H1-1A.3.2 (28 downstream strains).

**Table 2.**
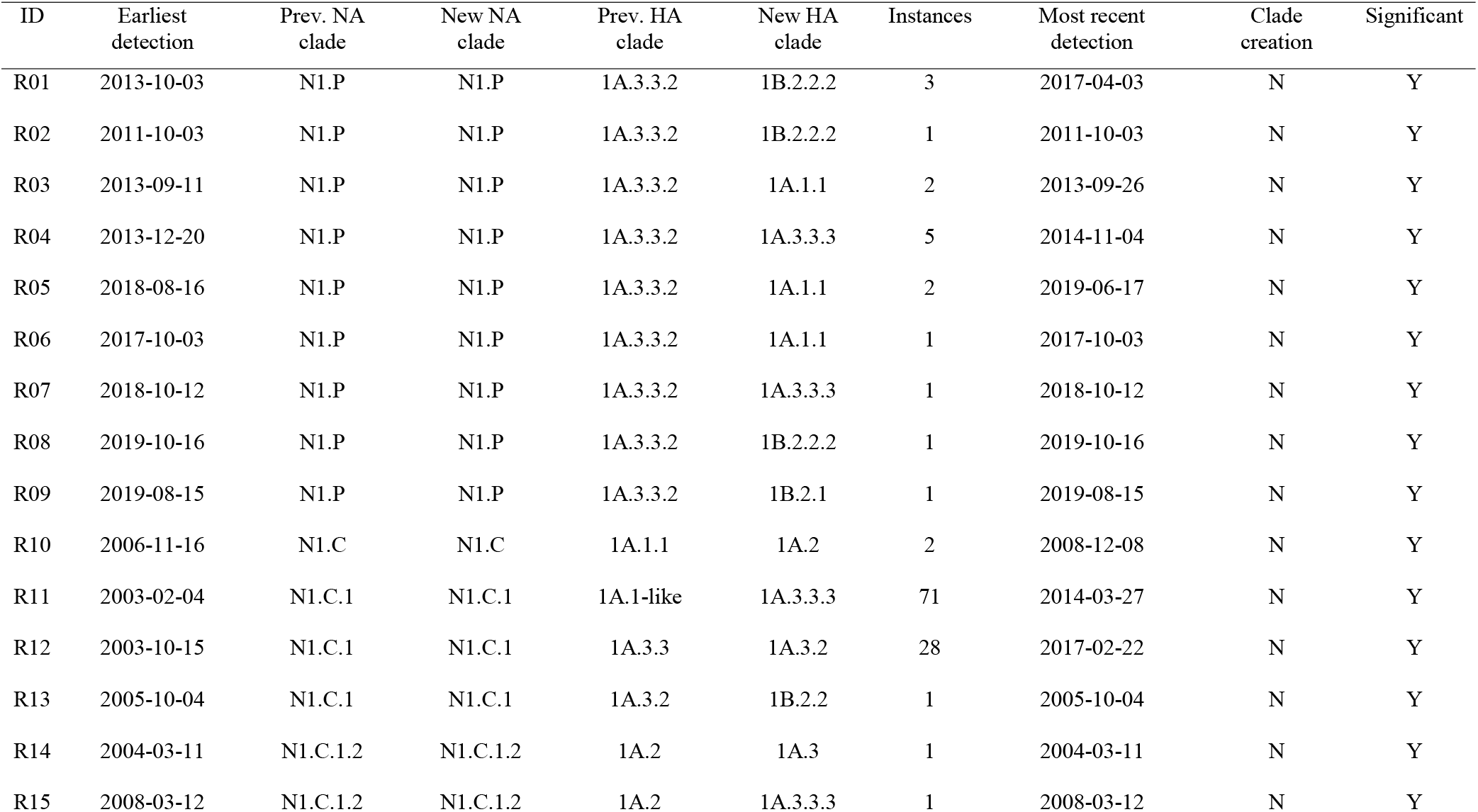
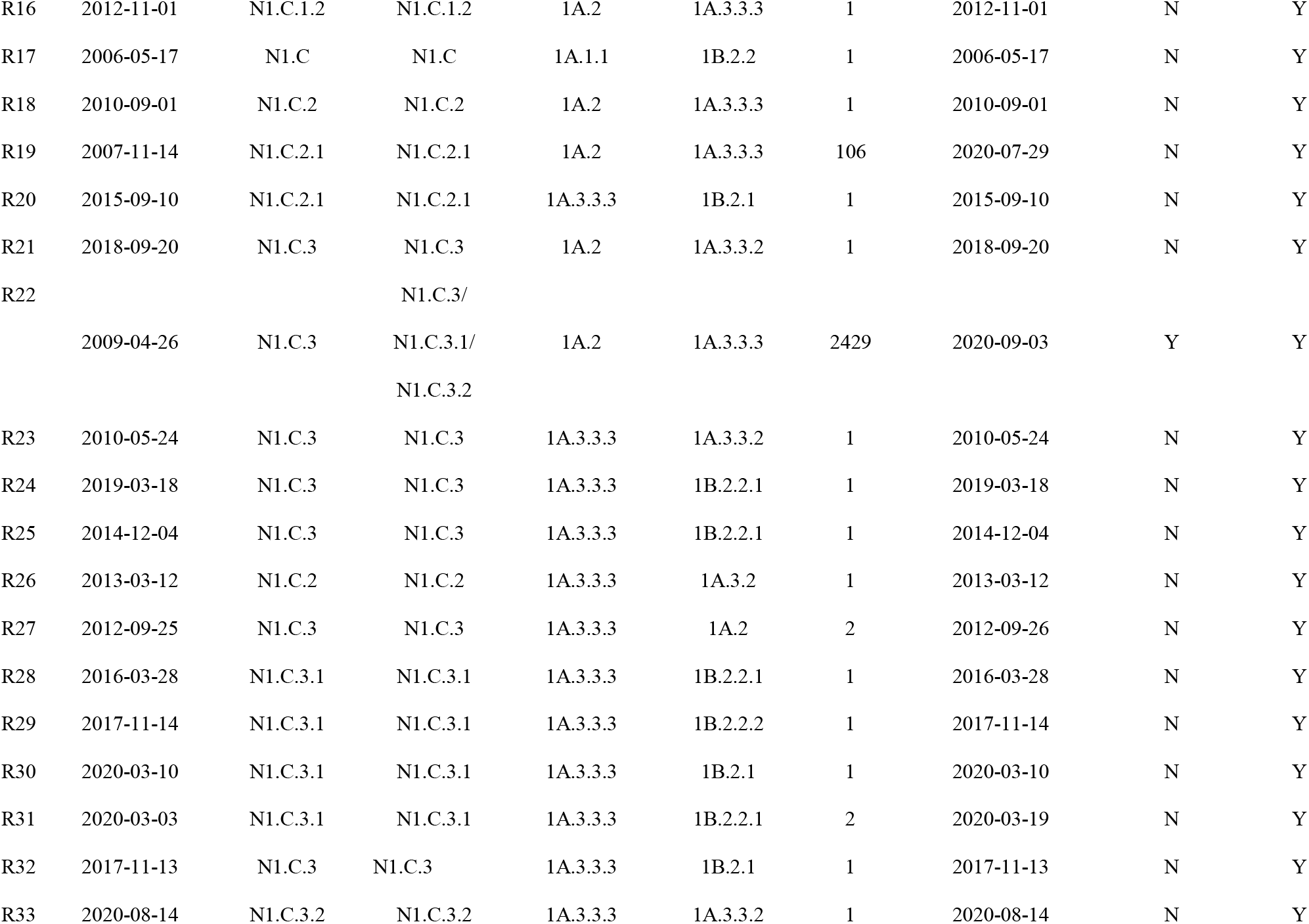
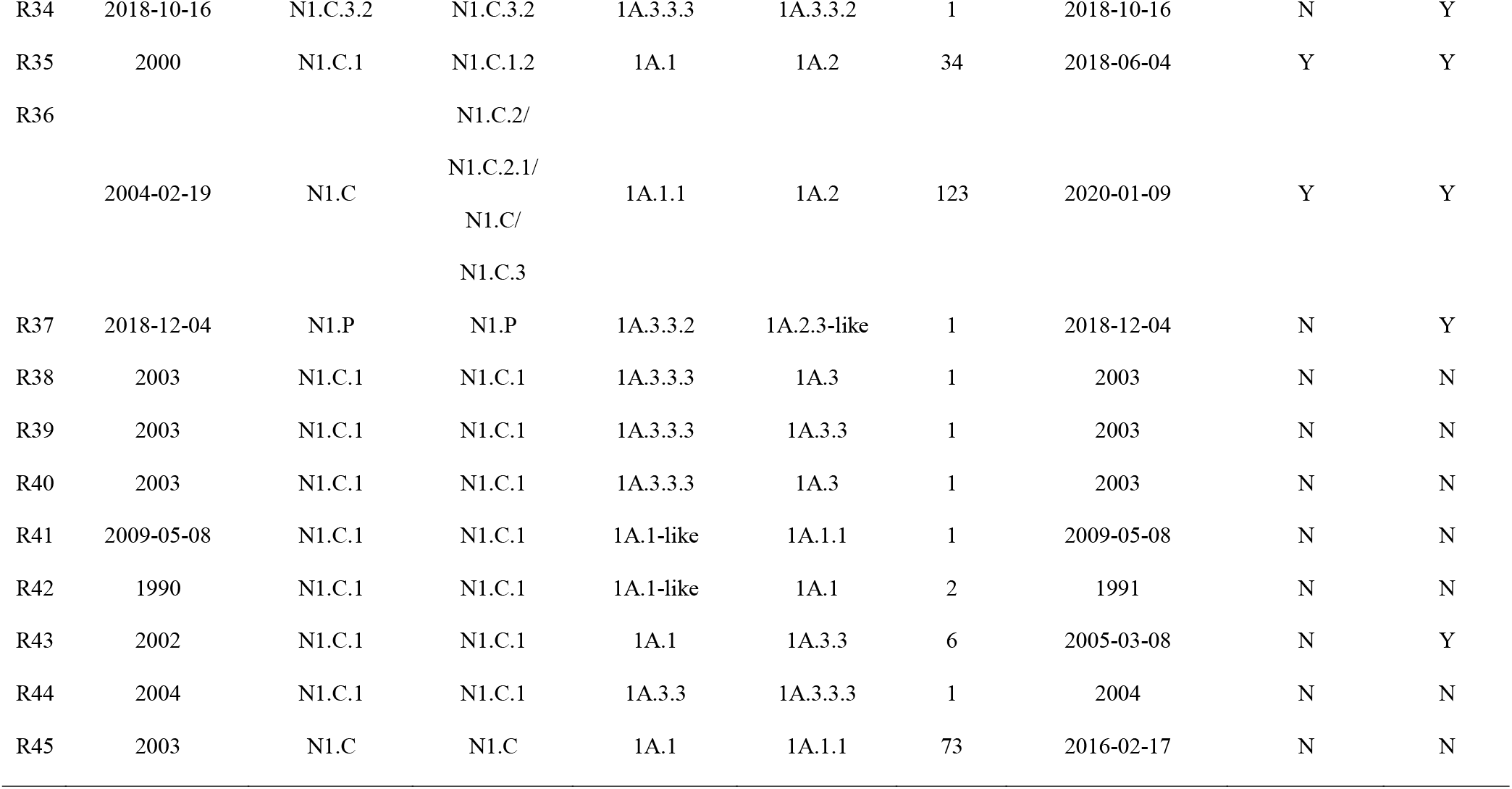
N1 neuraminidase reassortment events detected from influenza A virus in swine genetic clades. a) ongoing clades are clades with a detection in 2019 or more recently. b) a clade-defining reassortment event is an event that occurs at the start of a clade and includes all or most members of that clade

## 4. DISCUSSION

In this study we identified fourteen genetically distinct N1 NA clades within the Eurasian avian (EA) and classical swine (C) evolutionary lineages from swine in North America. Seven of the clades cocirculated between 2018 and 2020. The C lineage circulated in swine for a century (Worobey et al. 2014) and for most of this time it was genetically stable. In the mid-1980s the C lineage diverged, and new N1 clades have been detected subsequently at an increasing rate with an average of a new clade every 4 years. When a novel N1 clade emerged, it was associated with reassortment between N1 and HA. Between 2010 and 2020, viruses with specific N1 and N1-HA pairings moved throughout the US, creating transient hotspots of genetic diversity in single or multi-state clusters, usually lasting for one to two years at a time. The extent of the N1 genetic diversity suggests high adaptive potential, and this was supported by frequent detection of N1-HA reassortment. However, only a small fraction of the new N1-HA pairings persisted, plausibly because of intergene epistasis and functional balance. There was significant antigenic distance between the C and EA lineages, and there was a range of antigenic variation within the C lineage. These data suggest that there may be limited NA-mediated antibody immunity in swine to circulating N1 proteins of different clades. Our analysis shows how the adaptive potential of IAV in swine is mediated by coevolutionary dynamics of the surface proteins. As IAV fitness is largely determined by the ability to escape the host immune system, our work identifies how N1 diversity and N1-HA pairings resulted in significant antigenic variation that poses a challenge for controlling IAV in swine. We previously demonstrated dramatic antigenic diversity in swine H1 (Rajao et al. 2018) and here we show compounding antigenic diversity in the paired N1.

We found that preferential pairing and purifying selection are the major forces in the evolution of the N1 in swine. We identified six reassortment events across ninety-years that were sustained and changed the genetic landscape of the N1 in swine. For the clades N1.C.3, N1.C.1.2, and N1.C.2, the reassortment events R22, R35, and R36 were associated with the emergence of new clades and the newly acquired HA remained in the pairing across >99% of all subsequent detections. The number of reassortment events that failed to persist suggests that there are strong selective forces acting on N1-HA pairings, and this aligns with prior findings demonstrating the codependence of the HA and NA (Neverov et al. 2015, Gaymard et al. 2016). Though reassortment can lead to a temporary increase in the mutation rates in HA and NA (Ward et al. 2013, Neverov et al. 2015), and the NA specifically (Neverov et al. 2014), most novel reassorted N1-HA pairings were not maintained in the field. A concordant line of evidence is our data demonstrating purifying selection in all lineages and clades. Across the whole N1 gene there was a pattern for the third codon position to evolve faster than the first and second codon positions. There are likely to be many amino acid sites that function in coordination with the HA to attach and release from host cells that must be conserved to maintain biological function. While individual amino acids may be experiencing positive selection, if escape from the host immune response and positive selective pressure were the primary determinant of N1 gene evolution, the 1^st^ and 2^nd^ codon positions would have an increased gene-wide evolutionary rate relative to the 3^rd^ codon position. Instead our analyses show how diverse N1 genes reassort, with a small fraction of the resultant N1-HA pairings persisting with subsequent transmission and evolution that is likely constrained by the functional balance between the NA and HA genes.

We observed variation in the evolutionary rates of the N1 clades within and between lineages. Prior literature suggests that N1 evolves slower than N2 in swine (Jang and Bae 2018) and both NA subtypes evolve slower or at a comparable rate to the HA (Bhatt et al. 2013, Joseph et al. 2017, Jang and Bae 2018, Zeller et al. 2021). In our study, we demonstrated that the N1.P lineage had a significantly increased evolutionary rate relative to the C lineage. This elevated evolutionary rate for pdm09-derived sequences has also been seen in human strains, where the pdm09 NA evolved faster than NA genes from other human IAV (Xu et al. 2012). The N1.P gene in swine IAV reflects evolution in humans, multiple human-to-swine spillovers, and subsequent reassortment in swine (Nelson et al. 2015, Walia et al. 2019). Considering N1.P detections of the clade were heavily influenced by reverse-zoonosis, the rapid rate of evolution in swine may be due to a trait of the sequence itself or a positive selective pressure to adapt to a new host environment. Our results with N1.P are comparable to global EA lineage swine data that excluded the N1.P clade (Xu et al. 2012, Bhatt et al. 2013).

Including N1 clade-specific vaccine antigens in swine IAV vaccines that reflect the dominant N1 clades and N1-HA pairs in the US will likely improve vaccine efficacy. NI antibodies led to a reduction in both IAV infection and illness, independent of HA antibodies (Schulman 1969, Murphy et al. 1972, Monto and Kendal 1973, Couch et al. 2013). NI antibody titers have also been correlated with the efficacy of various vaccines (Clements et al. 1986, Monto et al. 2015). Vaccination for IAV in swine is a challenging problem given a rapidly evolving HA across more than 10 antigenically distinct H1 and H3 clades (Anderson et al. 2021). If the vaccine strain and circulating field strain HA are too antigenically dissimilar, vaccine induced antibody immunity may fail, but including an NA may broaden protection (Schulman 1969, Easterbrook et al. 2012, Rockman et al. 2013, Wan et al. 2013, Liu et al. 2015, Wohlbold et al. 2015) and reduce the potential for vaccine associated enhanced respiratory disease (Rajao et al. 2016, Wymore Brand et al. 2022). Our data demonstrate significant antigenic differences between the N1.P and C lineages, and certain N1 genetic clades were almost exclusively paired with specific HA genetic clades. Within the C lineage there was 2.0 AU variation across the 6 measured genetic clades, with each of the representative strains having at least one pairwise distance measure of greater than 3.0 AU. This suggests that a single C lineage gene may not be adequate to cover all N1 strains infecting North American swine populations. However, the major C lineage clades N1.C.3, N1.C.3.1 and N1.C.3.2 represented nearly 2500 genes and are approximately one third of IAV detected in our data set. Consequently, using surveillance to track dominant NA clades in the region of interest and using them in vaccines will improve average vaccine efficacy especially when swine are exposed to strains with a heterologous HA.

Our spatial analysis demonstrated that clade-specific N1 viruses moved throughout the country in spatial groups causing hotspots of infection, with each hotspot of N1 and N1-HA lasting one to two years. This temporal turnover in N1 diversity may partially reflect vaccine driven immunity. Since there was strong preferential pairing between N1-HA reflected in mirrored patterns in the N1 and N1-HA (Figure 5), as vaccine components are updated to reflect trends in circulating HA diversity, whole inactivated virus vaccines would concurrently include the paired N1 genes. There is also extensive use of custom or autogenous vaccines within swine in production systems to reflect on-farm surveillance activities. Consequently, as vaccine antigens are changed to reflect HA diversity, they are also likely targeting a different N1 genes. The swine immune profile then could provide an advantage to specific N1 and N1-HA genes that we detected as one-to-two-year spatiotemporal clusters (Figure 5).

The pace of diversification for the N1 NA increased with time, but this increased diversity is occurring in the context of a long evolutionary history; the consequence of which was a limited number of N1-HA pairings and less antigenic diversity compared to N2 in swine (Kaplan et al. 2021). A likely contributor is that N1 is nearly always paired with an H1 whereas N2 pair with both H1 and H3 in swine IAV (Arendsee et al. 2021) and may therefore be less affected by epistatic pressure. The C lineage appeared to have little genetic or antigenic change for seventy years (Olsen 2002, Vincent et al. 2006, Nelson et al. 2011), with a single genetic clade consistently paired with a single H1. Major evolutionary events such as the detection and spread of the triple reassorted H3N2 viruses in ~1998 (Vincent et al. 2006, Vincent et al. 2020), and the H1 1B human seasonal lineage in the early 2000s (Vincent et al. 2009), appear to have done little to change the preferential pairings seen in C lineage N1 genes. Our data demonstrated that the genetic landscape is dynamic, with new N1 genetic clades being detected approximately every 4 years. The rate of detection of N1 IAV clades increased in frequency, concurrent with reverse zoonoses and reassortment associated with the H1N1pdm09 (Nelson et al. 2015, Gao et al. 2017). Reassortment was associated with genetic and antigenic diversity and may have increased the rate at which new N1-HA pairings emerged across North America. Based on the evidence we present on preferential pairing, purifying selection, and reduced evolutionary rate concurrent with substantial N1 genetic diversity, these data suggest that N1 genes overall reached an equilibrium near the peak of virus fitness in the swine host.

## 5. CONCLUSION

In this project we quantified the genetic and antigenic diversity of N1 NA in North American swine IAV. This study demonstrated seven currently circulating genetic clades, unique antigenic phenotypes of clades with substantial differences between lineages, diverse rates of evolution, highly variable spatial and temporal patterns in detection frequency in genes and HA-N1 gene pairings and demonstrated how rare reassortment events shaped the evolutionary history of N1 genes in swine. We identified purifying selection and preferential pairing as meaningful forces acting upon the N1 NA, perhaps due to a long and stable evolutionary history that led to a fitness peak. The classification system for swine N1 established here facilitates communication about strains that are in circulation and provides a baseline to detect clades as they expand or contract in diversity, reassort, or emerge in distinct geographic areas.

## ACKNOWLEDGEMENTS

We gratefully acknowledge pork producers, swine veterinarians, and laboratories for participating in the USDA Influenza A Virus in Swine Surveillance System and publicly sharing sequences.

## FUNDING

This work was supported in part by: the U.S. Department of Agriculture (USDA) Agricultural Research Service [ARS project number 5030-32000-231-000-D; ARS project number 5030-32000-231-025-S]; the National Institute of Allergy and Infectious Diseases, National Institutes of Health, Department of Health and Human Services [contract number 75N93021C00015 and 75N93021C00014]; the USDA Agricultural Research Service Research Participation Program of the Oak Ridge Institute for Science and Education (ORISE) through an interagency agreement between the U.S. Department of Energy (DOE) and USDA Agricultural Research Service [contract number DE-AC05-06OR23100]; the Department of Defense, Defense Advanced Research Projects Agency, Preventing Emerging Pathogenic Threats program [contract number HR00112020034]; and the SCINet project of the USDA Agricultural Research Service [ARS project number 0500-00093-001-00-D]. The funders had no role in study design, data collection and interpretation, or the decision to submit the work for publication. Mention of trade names or commercial products in this article is solely for the purpose of providing specific information and does not imply recommendation or endorsement by the USDA, DOE, or ORISE. USDA is an equal opportunity provider and employer.

## DATA AVAILABILITY

The code and data in this manuscript are provided at https://github.com/flu-crew/N1-diversity or at USDA Ag Data Commons doi: https://doi.org/10.15482/USDA.ADC/1528285 Supplementary figures are available online at XXX.

## Notes

https://github.com/flu-crew/N1-diversity

https://doi.org/10.15482/USDA.ADC/1528285

### Competing Interest Statement

The authors have declared no competing interest.

