## Supplemental Material for "Characterizing a century of genetic diversity and contemporary antigenic diversity of N1 neuraminidase in IAV from North American swine"

### Supplementary Figures and Tables

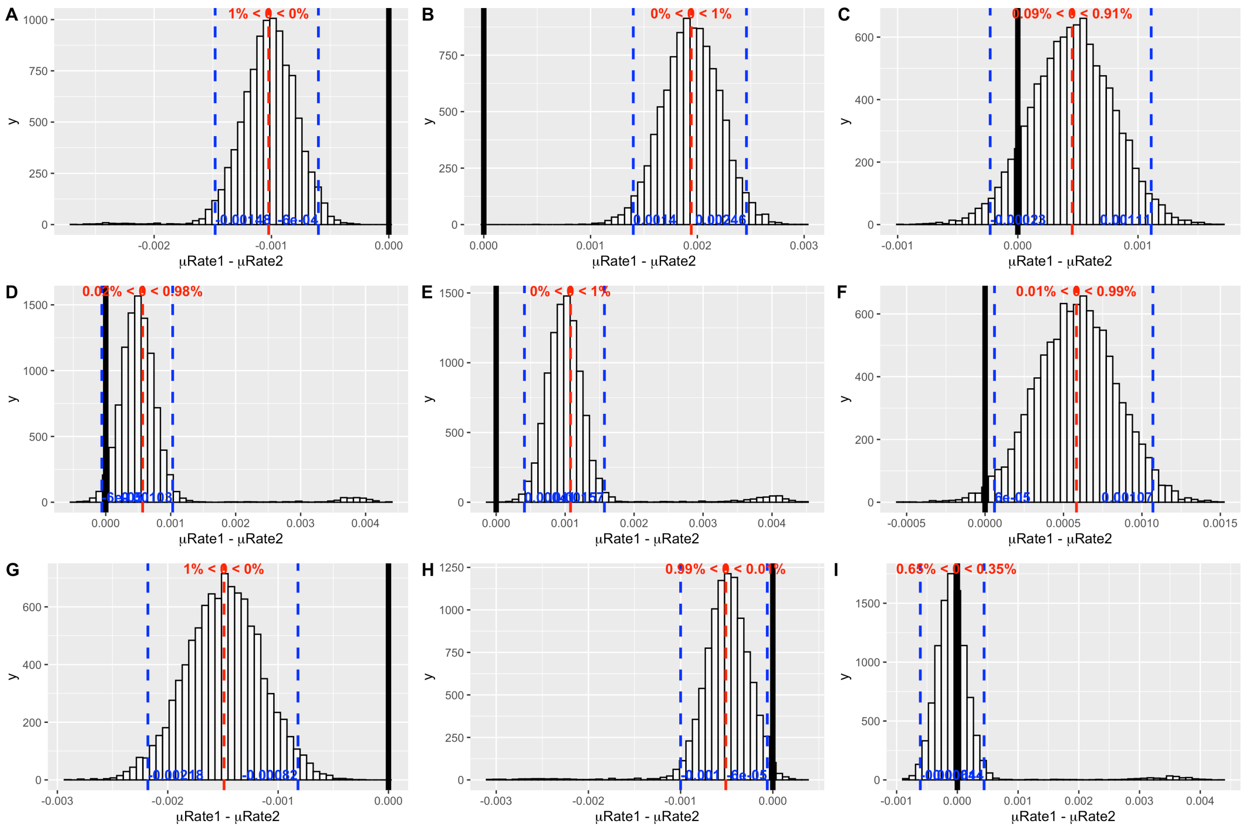


**Figure S1**. Plots of one posterior distribution of substitution rates from BEAST minus another for A) N1.C vs C. lineage, B) N1.P vs N1.C.2, C) N1.P vs N1.C.2.1, D) N1.P vs. N1.C.3, E) N1.P vs. N1.C.3.1, F) N1.P vs. N1.C.3.2, G) N1.C.2.1 vs N1.C.2, H) N1.C.3 vs N1.C.3.1, I) N1.C.3 vs. N1.C.3.2. Vertical blue lines indicate the credible interval, vertical red lines indicate the mean of the subtracted distributions, and the vertical black line indicates zero.

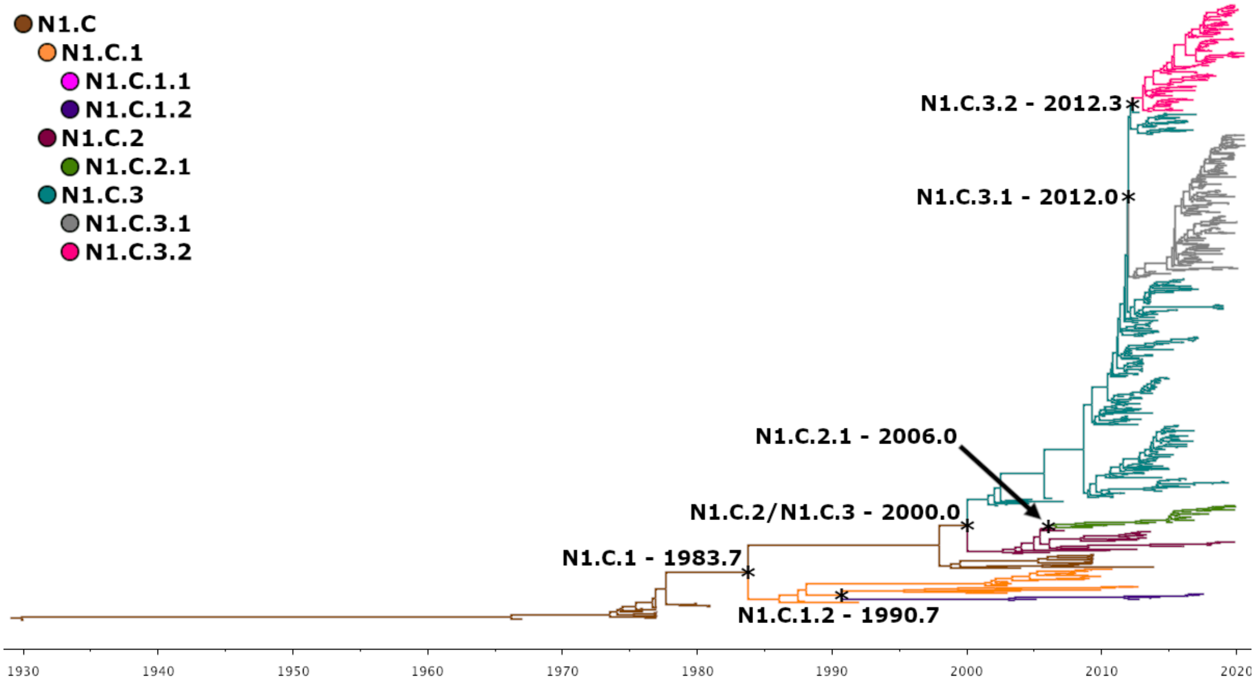


**Figure S2.** A time-scaled maximum clade credibility tree of the N1 neuraminidase influenza A virus classical swine lineage. Stars indicate internal nodes where new clades diverged. These clades are named and the time to the most recent common ancestor are displayed.

Table S1: Clade distribution across regions and in our data set including summaries of most prevalent regions, and all other regions where clades were found

| clade | Number clade | Percent clade in data | Major region | number clade in region | Percent clade in region | Percent region in all data | Other region |
|---|---|---|---|---|---|---|---|
| N1.H | 11 | 0.29 | NC | 5 | 45.45 | 11.53 | CAN,NC,IL,MN,SC |
| N1.A | 1 | 0.03 | NA | NA | NA | NA | CAN |
| N1.E | 1 | 0.03 | NA | NA | NA | NA | MEX |
| N1.E.1 | 18 | 0.48 | MEX | 18 | 100.00 | 1.94 | MEX |
| N1.P | 629 | 16.86 | IA | 111 | 17.65 | 29.14 | CAN,MEX,CRI,GTM,CUB,NC,OK,VA,AR,TX,WI,KS,UT,NE,TN,GA,PA,KY,IL,IN,IA,CO,OH,MO,MN,OR,SD |
| N1.C | 171 | 4.58 | CAN | 76 | 44.44 | 5.19 | CAN,NJ,AZ,WI,CO,OH,OK,MN,NE,KY,TN,IA,IL |
| N1.C.1 | 125 | 3.35 | NC | 43 | 34.40 | 11.53 | CAN,MEX,OR,NC,WI,OH,MO,MN,KY,OK,TN,KS,GA,CA,IL,IN,IA,MD,SD,TX |
| N1.C.1.1 | 22 | 0.59 | IA | 5 | 22.73 | 29.14 | OH,OK,MN,NE,CA,IL,IN,IA,TX,SD |
| N1.C.1.2 | 37 | 0.99 | CAN | 10 | 27.03 | 5.19 | CAN,NC,PA,MN,VA,KY,TN,AR,IL,IA,SD |
| N1.C.2 | 109 | 2.92 | NE | 36 | 33.03 | 3.63 | MEX,MN,CO,OH,OK,KS,NE,AR,IL,IN,IA,MO,TX,SD |
| N1.C.2.1 | 122 | 3.27 | NC | 90 | 73.77 | 11.53 | MS,MN,OH,MO,NC,UT,VA,IL,IN,IA,TX |
| N1.C.3 | 1230 | 32.98 | IA | 269 | 21.87 | 29.14 | MS,NC,OK,VA,AR,TX,AZ,WI,KS,NE,CA,PA,KY,ND,IN,IA,CO,OH,MO,MN,MI,SC,SD,IL |
| N1.C.3.1 | 711 | 19.06 | IA | 332 | 46.69 | 29.14 | KS,OK,VA,AR,TX,WI,NC,NE,TN,CA,PA,KY,IL,IN,IA,CO,OH,MO,MN,MI,SC,SD |
| N1.C.3.2 | 543 | 14.56 | IA | 324 | 59.67 | 29.14 | NC,OH,PA,MI,MO,MN,FL,NE,TN,KS,AR,IL,IN,IA,AL,OK,SD |

Table S2. Representative N1 neuraminidase and paired hemagglutinin genes from contemporary circulating clades of N1 in North America. The N1 gene was used to generate recombinant H9N1 for use in neuraminidase inhibition enzyme-linked lectin assays (NI ELLA).

| **Strain name** | **N1 clade** | **GenBank Accession** | **HA clade** | **GenBank Accession** |
|---|---|---|---|---|
| A/swine/Iowa/A02479002/2020 | N1.P | MT052095 | 1A.3.3.2 | MT052094 |
| A/swine/California/A02478680/2019 | N1.C.1.1 | MN575727 | 1A.2-3-like | MN575726 |
| A/swine/South Dakota/A02016893/2018 | N1.C.1.2 | MH551255 | 1A.2 | MH551254 |
| A/swine/Kansas/A02245346/2019 | N1.C.2 | MN857569 | 1A.2 | MN857568 |
| A/swine/North Carolina/A02479062/2020 | N1.C.2.1 | MT154208 | 1A.3.3.3 | MT154207 |
| A/swine/Minnesota/A02245409/2020 | N1.C.3.1 | MT154190 | 1A.3.3.3 | MT154189 |
| A/swine/Iowa/A02479110/2020 | N1.C.3.2 | MT277490 | 1A.3.3.3 | MT277489 |

Table S3: Available molecular substitution rates for converged and best replicate lineages and clades with at least 50 sequences and evidence of ongoing circulation

| Lineage/clade | Sample size | Subsampled? | Likelihood | Substitution rate[a] | 1,2, sub. rate[b] | 3 sub. rate[c] |
|---|---|---|---|---|---|---|
| Classical lineage | 3064 | Y (n=750) | -30634.3 | 3.69E-03 | 2.07E-03 | 6.94E-03 |
| N1.C.2 | 109 | N | -6167.2 | 2.76E-03 | 1.27E-03 | 5.74E-03 |
| N1.C.2.1 | 122 | N | -5650.4 | 4.25E-03 | 2.27E-03 | 8.20E-03 |
| N1.C.3 | 1230 | Y (n=750) | -19641.9 | 4.22E-03 | 2.51E-03 | 7.65E-03 |
| N1.C.3.1 | 711 | N | -14511.8 | 3.73E-03 | 1.98E-03 | 7.22E-03 |
| N1.C.3.2 | 543 | N | -11980.8 | 4.12E-03 | 2.25E-03 | 7.86E-03 |
| N1.P | 629 | N | -18897.1 | 4.72E-03 | 2.90E-03 | 8.36E-03 |

[a]the general molecular substitution rate
[b]the absolute molecular substitution rate for the first and second base pairs in each codon
[c]the absolute molecular substitution rate for the third base pair in each codon

Table S4: Statistics and dates related to clade divergence and oldest and most recent sequences detected

| Clade | Divergence time[a] | Oldest strain | Oldest strain (decimal)[b] | Newest strain | Newest strain (decimal) | Years until detection[c] | Years in circulation[d] |
|---|---|---|---|---|---|---|---|
| N1.C | 1929.8 | 1930 | NA | 2/17/16 | 2016.1 | NA | 86.3 |
| N1.C.1 | 1983.7 | 1987 | NA | 2/22/17 | 2017.1 | 3.30 | 33.4 |
| N1.C.1.1 | NA | 1999 | NA | 11/12/19 | 2019.9 | NA | >=19.9 |
| N1.C.1.2 | 1990.7 | 2000 | NA | 6/4/18 | 2018.5 | 9.30 | 27.8 |
| N1.C.2 | 2000.0 | 2/19/04 | 2004.13 | 1/9/20 | 2020.0 | 4.13 | 20.0 |
| N1.C.2.1 | 2006.0 | 4/25/07 | 2007.32 | 7/29/20 | 2020.6 | 1.32 | 14.6 |
| N1.C.3 | 2000.0 | 5/16/02 | 2002.37 | 12/31/19 | 2020.0 | 2.37 | 20.0 |
| N1.C.3.1 | 2012.0 | 7/27/12 | 2012.57 | 8/27/20 | 2020.7 | 0.57 | 8.7 |
| N1.C.3.2 | 2012.3 | 8/27/13 | 2013.65 | 9/3/20 | 2020.7 | 1.35 | 8.4 |
| N1.H | NA | 2005 | NA | 7/20/09 | 2009.6 | NA | 3.6 |
| N1.A | NA | 2002 | NA | 2002 | NA | NA | NA |
| N1.E | NA | 1/8/13 | 2013.02 | 1/8/13 | 2013.0 | NA | NA |
| N1.E.1 | NA | 3/25/10 | 2010.23 | 9/30/15 | 2015.8 | NA | >=5.5 |
| N1.P | NA | May-09 | 2009.37 | 8/21/20 | 2020.6 | NA | >=11.3 |

[a]determined by BEAST-derived MCC tree except for N1.C where it is derived from the age statistic in a BEAST log file

[b]All decimal calculations are not performed when month is missing and when year and month only are present the day is assumed to be the 15th

[c]Years until detection is minimized when the month is not known for the oldest sequence

[d]Years in circulation is calculated based on divergence time when available. When unavailable earliest sequence is used and the ">=" signifier is used. Whenever only year is available for the earliest sequence the end of the year is used.
